# A Proteomics-Based Comparison of Host Responses to Spotted Fever Group *Rickettsia* in Endothelial Cells

**DOI:** 10.1101/2025.07.24.666595

**Authors:** Pedro Curto, Catia Santa, Bruno Manadas, Isaura Simoes

**Affiliations:** I3S - Instituto de Investigação e Inovação em Saúde, Porto, Portugal; IBMC - Instituto de Biologia Molecular e Celular, Universidade do Porto, Portugal; CNC-UC - Center for Neuroscience and Cell Biology, University of Coimbra; CiBB - Centre for Innovative Biomedicine and Biotechnology, University of Coimbra; Department of Biological Sciences, Old Dominion University, Norfolk, Virginia, USA

**Keywords:** Spotted fever group *Rickettsia*, *Rickettsia africae*, *Rickettsia parkeri*, *Rickettsia massiliae*, *R. montanensis*, SWATH-MS/MS, endothelial cells, HUVEC/TERT2, antiviral immunity, Type I interferon signaling, THP-1

## Abstract

Spotted fever group (SFG) *Rickettsia* species are obligate intracellular bacteria with a tropism for endothelial cells (ECs), where they initiate pathogenesis leading to rickettsial vasculitis. However, how endothelial cells sense and respond to infection by *Rickettsia* species of differing pathogenic potential remains poorly defined. In this study, we conducted a comparative analysis of four SFG *Rickettsia* species - *R. africae*, *R. parkeri*, *R. massiliae*, and *R. montanensis* - using high-throughput label-free SWATH/DIA-MS/MS in human HUVEC/TERT2 cells. Our results revealed distinct intracellular growth dynamics that correlated with known virulence profiles: the more pathogenic *R. africae* and *R. parkeri* replicated more efficiently, while the non-pathogenic *R. montanensis* failed to replicate. Proteomic profiling uncovered both shared and species-specific host responses, with a marked induction of proteins associated with type I interferon (IFN-I) signaling, particularly in response to *R. africae* and *R. parkeri*. Proteins typically involved in antiviral immunity, such as RIG-I, ISG15, IFITs, MX1, MX2, and OAS family members, were significantly accumulated, suggesting activation of cytosolic nucleic acid sensing pathways upon infection with pathogenic rickettsiae. ISGylation levels, however, remained low and varied depending on the species, pointing to complex regulatory mechanisms. Comparison with previous quantitative proteomics data in THP-1 macrophages revealed a conserved interferon signature, while also highlighting cell-type-specific responses. Overall, our findings demonstrate that endothelial cells activate innate immune pathways typically associated with antiviral defense upon *Rickettsia* infection. These immune signatures may serve as potential indicators of pathogenic potential and provide a foundation for identifying biomarkers and therapeutic targets in rickettsial diseases.

## INTRODUCTION

Vector-borne diseases are gaining global momentum due to economic globalization, changes in land use and urbanization, increases in travel, and global warming, all of which have been postulated to raise the distribution and incidence of these diseases [1–6]. Among these pathogens are different species of *Rickettsia*, causing severe infections like epidemic typhus (*R. prowazekii*), Rocky Mountain Spotted Fever (RMSF, *R. rickettsii*), and Mediterranean Spotted Fever (MSF, *R. conorii*) [7, 8]. Rickettsial infections, except for *R. akari*, are characterized by their affinity to preferentially infect vascular endothelial cells (ECs) lining the small and medium-sized blood vessels in humans and also in animal models of infection [8–11]. As a consequence, rickettsiae can disseminate through the endothelium, damaging vascular networks, which leads to disseminated inflammation, loss of barrier function, and altered vascular permeability (collectively referred to as rickettsial vasculitis) [12, 13]. Indeed, most of the clinical features of rickettsial diseases have been attributed to disseminated infection of the endothelium, where *Rickettsia* cause oxidative stress, thereby causing injury to the endothelial cells [8, 14, 15]. However, during rickettsial infections, ECs are not merely injured but are also able to launch an array of adaptive cellular responses, switching from a basal and non-thrombogenic phenotype to a state known as “endothelial activation” [12, 13, 16]. The responses that characterize an “activated endothelial” state have been the subject of several studies, and hallmark features include, but are certainly not limited to, higher expression of pro-thrombotic, pro-adhesive, and pro-inflammatory genes [8, 13, 16]. Although chemokines are expressed at relatively low levels in endothelial cells, increased levels of interleukins and chemokines have been reported upon infection with rickettsiae [13]. The production of inflammatory cytokines, such as IL-1α, IL-6, and IL-8 by rickettsiae-infected endothelial cells has been correlated with the expression of cell adhesion molecules, such as intercellular-adhesion molecule 1 and vascular-cell-adhesion molecule 1, which support the recruitment of T cells to the site of infection. Increased expression of CCL2, CCL3, CCL4, and CCL5 has also been correlated with macrophage and monocyte interactions with endothelial cells, while increased levels of CXCL5 and CXCL8 are reported to act in recruitment of monocytes, macrophages, lymphocytes, and other polymorphonuclear leukocytes to the site of infection, and CXCL9 and CXCL10 as T-cell chemoattractants [8, 10, 17–20]. The peak of expression of chemokines also correlates with maximal T-cell infiltration (mainly CD8+ T cells) at the site of infection, suggesting a regulatory T-cell and endothelial cell crosstalk.

In fact, ECs have emerged as key immunoreactive cells that participate in a diverse array of cellular processes by both producing and/or reacting to a broad range of mediators, being considered a key cell type during innate and adaptive immunity [21–23]. Due to its constant and close contact with the bloodstream, ECs serve as sentinels by sensing invading pathogens. High-throughput proteomics approaches were already employed to study the endothelial cell responses to *R. conorii* infection, revealing an increase in IFN and MHC class I antigen presentation pathways [24]. However, it remains elusive how endothelial cells sense infection signals and mount an effective immune response during rickettsial infections

To gain deeper insights into the endothelial cell responses to infection with different spotted fever group (SFG) *Rickettsia* species with diverse degrees of pathogenicity to humans, we infected the hTERT-immortalized human umbilical vein endothelial (HUVEC/TERT2) cell line with *R. africae*, *R. parkeri*, *R. massiliae*, and *R. montanensis*, and assessed bacterial replication dynamics by immunofluorescence microscopy. Our results showed that SFG *Rickettsia* species display distinct growth kinetics in HUVEC/TERT2 cells, with the more pathogenic species replicating more efficiently. To understand the host cellular responses underlying these differences, we applied an unbiased, label-free high-throughput proteomic approach (SWATH/DIA-MS/MS) to globally profile endothelial responses to each species. Infection with the more pathogenic *Rickettsia* species was associated with the increased accumulation of proteins involved in innate immune and type I interferon responses, including the cytosolic RNA sensor RIG-I, the ubiquitin-like modifier ISG15, IFIT family members, and the antiviral GTPases MX1 and MX2. We also observed increased levels of oligoadenylate synthetases (OALS, OAS2, and OAS3), further supporting the activation of interferon-stimulated antiviral pathways. These data suggest that endothelial cells trigger antiviral sensing pathways in response to *Rickettsia* infection, and that the magnitude and nature of these responses likely reflect the pathogenic potential of different SFG *Rickettsia* species.

## MATERIALS AND METHODS

### Cell lines and *Rickettsia* growth and purification

Vero cells were grown in Dulbecco’s modified Eagle’s medium (DMEM; Gibco) supplemented with 10% heat-inactivated fetal bovine serum (Atlanta Biologicals), 1× nonessential amino acids (Corning), and 0.5 mM sodium pyruvate (Corning). HUVEC/TERT2 (CHT-006-0008; Evercyte) were grown in Endothelial Basal Medium (EBM; Lonza) supplemented with EGM™ Endothelial Cell Growth Medium SingleQuots™ Kit (Lonza). For cell culture of HUVEC/TERT2, cell culture flasks/plates were pre-treated with a solution of 0.1% gelatin in PBS for at least 10 minutes. Both cell lines were maintained in a humidified 5% CO_2_ incubator at 34°C. *Rickettsia montanensis* isolate M/5-6T, *Rickettsia massiliae* isolate MTU5, *Rickettsia parkeri* isolate Portsmouth, and *Rickettsia africae* isolate ESF-5 T were obtained from CSUR-Collection de Souches de l’Unité des Rickettsies, Marseille, France. All *Rickettsia* species were propagated in Vero cells and purified as previously described [25, 26].

### Assessment of *Rickettsia* growth dynamics

Growth dynamics was assessed by inoculating *R. parkeri*, *R. africae*, *R. massiliae*, and *R. montanensis* at a multiplicity of infection (MOI) of 10 into HUVEC/TERT2 cells at a confluency of 2 × 10^5^ cells per well, in 24-well plates seeded onto glass coverslips. Plates were centrifuged at 300 × *g* for 5 min at room temperature to induce contact between rickettsiae and host cells and incubated at 34°C and 5% CO_2_. At each specific time point post-inoculation, infected monolayers were washed with PBS and fixed in 4% paraformaldehyde (PFA) for 20 min. Samples were then permeabilized with 0.1% Triton X-100 and blocked with 2% bovine serum albumin (BSA). *Rickettsia* growth was assessed by staining with anti-*Rickettsia* polyclonal antibody NIH/RML I7198 (1:1,500), followed by Alexa Fluor 488-conjugated goat anti-rabbit IgG (1:1,000), DAPI (4′,6-diamidino-2-phenylindole; 1:1,000), and Texas Red–X phalloidin (1:200). After washing with PBS, glass coverslips were mounted in Mowiol mounting medium and preparations were viewed on a Zeiss Axiovert 200M fluorescence microscope (Carl Zeiss) using a final 40× optical zoom and processed with ImageJ software.

### Mass spectrometry sample preparation

HUVEC/TERT2 cell monolayers at a cell confluence of 2 × 10^5^ cells per well, in 24-well plates (6 wells per condition), were infected with *R. parkeri*, *R. africae*, *R. massiliae*, and *R. montanensis* at an MOI of 10 or maintained uninfected as a control. Plates were centrifuged at 300 × *g* for 5 min at room temperature to induce contact between rickettsiae and host cells and incubated at 34°C and 5% CO2 for 24 h. At the specified time point, the culture medium was removed, cells were washed 1× with PBS, and total protein was extracted using 100 μL of protein extraction buffer per well (25 mM Tris/HCl, 5 mM EDTA, 1% Triton X-100, and Pierce protease inhibitors [Thermo Fisher Scientific], pH 7.0). Samples were passed 10 times through an insulin syringe with a 28-gauge needle (Becton, Dickinson) and denatured using 6× SDS sample buffer (4× Tris/HCl, 30% glycerol, 10% SDS, 0.6 M dithiothreitol, 0.012% bromophenol blue, pH 6.8) for 10 min at 95°C. The total protein content in each sample was then quantified using the Pierce 660nm protein assay kit (Thermo Fisher Scientific) and kept at −80°C until further processing. Experiments were done in quadruplicate. After thawing, amounts of 10 μg of each replicate sample from each experimental condition were pooled, creating the five pooled samples (*R. parkeri* pool, *R. africae* pool, *R. massiliae* pool, *R. montanensis* pool, and uninfected pool). At this point, the same amount of a recombinant protein (green fluorescent protein fused to maltose-binding periplasmic protein [MalE-GFP]) was added to each replicate sample and the pooled samples to serve as an internal standard. All the samples were boiled for 5 min, and acrylamide was added as an alkylating agent.

### In-gel digestion and LC-MS/MS

The volume corresponding to 40 μg of each replicate sample, as well as pooled samples, was then loaded into a precast gel (4 to 20% Mini-Protean TGX gel; Bio-Rad), and SDS-PAGE was partially run at 110 V [27]. After SDS-PAGE, proteins were stained with colloidal Coomassie blue as previously described [28]. The lanes were sliced into 3 fractions with a scalpel, and after the excision of the gel bands, each one was sliced into smaller pieces. The gel pieces were destained using a 50 mM ammonium bicarbonate solution with 30% acetonitrile (ACN), followed by a washing step with water (each step was performed in a thermomixer [Eppendorf] at 1,050 × rpm for 15 min). The gel pieces were dehydrated on a Concentrator plus/Vacufuge R plus (Eppendorf). To each gel band, 75 μL of trypsin (0.01 μg/μL solution in 10 mM ammonium bicarbonate) was added and the band left for 15 min at 4°C to rehydrate the gel. After this period, 75 μL of 10 mM ammonium bicarbonate was added and in-gel digestion was performed overnight at room temperature in the dark. After digestion, the excess solution from gel pieces was collected into a low-binding microcentrifuge tube (LoBind, Eppendorf) and peptides were extracted from the gel pieces by the sequential addition of three solutions of increasing percentages of ACN (30, 50, and 98%) in 1% formic acid (FA). After the addition of each solution, the gel pieces were shaken in a thermomixer (Eppendorf) at 1,250 rpm for 15 min and the solution was collected into the tube containing the previous fraction. The peptide mixtures were dried by rotary evaporation under vacuum (Concentrator plus/Vacufuge plus; Eppendorf). The peptides from each fraction of each sample were pooled for SWATH analysis, while the peptides from the pooled samples were kept separated in the three fractions of the digestion procedure. After digestion, all samples were subjected to solid-phase extraction with C18 sorbent (OMIX tip; Agilent Technologies). The eluted peptides were evaporated and solubilized in 30 μL mobile phase, aided by ultrasonication using a cup horn device (Vibra-Cell 750 W; Sonics) at 40% amplitude for 2 min. Samples were then centrifuged for 5 min at 14,100 × *g* (MiniSpin plus; Eppendorf) and analyzed by liquid chromatography coupled to tandem mass spectrometry (LC-MS/MS). The TripleTOF 6600 system (Sciex) was operated in two phases: Data-dependent acquisition (DDA) of each fraction of the pooled samples, followed by SWATH/DIA (sequential windowed data-independent acquisition of the total high-resolution mass spectra/Data Independent Acquisition) of each sample. Peptide separation was performed using liquid chromatography (nanoLC 425; Eksigent) on a Triart C18 capillary column 1/32 in (12 nm, S-3μm, 150 by 0.3 mm; YMC) and using a Triart C18 capillary guard column (0.5 by 5 mm, 3 μm, 12 nm; YMC) at 50°C at 5 μL/min with a 50-min gradient from 5% to 30% ACN in 0.1% FA in a total run time of 65 min, and the peptides were eluted into the mass spectrometer using an electrospray ionization source (DuoSpray source; Sciex). IDA experiments were performed by analyzing each fraction of the pooled samples. The mass spectrometer was set for DDA scanning full spectra (350 to 2,250 m/z) for 250 ms, followed by 50 MS/MS scans (100 to 1,500 m/z with an accumulation time of 60 ms to maintain a cycle time of 3.3 s). Candidate ions with a charge state between +1 and +5 and counts above a minimum threshold of 100 cps were isolated for fragmentation, and one MS/MS spectrum was collected before adding those ions to the exclusion list for 15 s (mass spectrometer operated by Analyst TF 1.8; Sciex). Rolling collision energy was used with a collision energy spread (CES) of 5. The SWATH/DIA setup was essentially as described previously [29], with the same chromatographic conditions used for SWATH and DDA acquisitions. For SWATH-MS-based experiments, the mass spectrometer was operated in a looped product ion mode. The SWATH-MS setup was designed specifically for the samples to be analyzed, in order to adapt the SWATH windows to the complexity of this batch of samples. A set of 168 windows of various widths (containing 1 m/z for window overlap) was constructed covering the precursor mass range of 350 to 2,250 m/z. A 50-ms survey scan (350 to 2,250 m/z) was acquired at the beginning of each cycle for instrument calibration, and SWATH-MS/MS spectra were collected from 100 to 1500 m/z for 19 ms, resulting in a cycle time of 3.76 s from the precursors ranging from 350 to 2,250 m/z. The collision energy for each window was determined according to the calculation for a charge +2 ion centered upon the window, with varying collision energy spread according to the window.

### Protein identification and relative quantification

A specific library of precursor masses and fragment ions was created by combining all files from the IDA experiments and used for subsequent SWATH processing. The library was obtained using Protein Pilot™ software (version 5.0.1; Sciex) with the following search parameters: Homo sapiens Swiss-Prot database (downloaded in September 2020) and MalE-GFP, acrylamide-alkylated cysteines as fixed modification, and the gel-based special focus option. An independent false discovery rate (FDR) analysis using the target-decoy approach provided with the Protein Pilot software was used to assess the quality of the identifications, and identifications were considered positive when identified proteins and peptides reached a 5% local FDR [30, 31]. Data processing was performed using the SWATH processing plug-in for PeakView™ (version 2.2; Sciex). Briefly, peptides were selected from the library using the following criteria: (i) the unique peptides for a specific targeted protein were ranked by the intensity of the precursor ion from the IDA analysis as estimated by the Protein Pilot software, and (ii) peptides that contained biological modifications and/or were shared between different protein entries/isoforms were excluded from selection. Up to 15 peptides were chosen per protein, and SWATH quantitation was attempted for all proteins in the library file that were identified below 5% local FDR from Protein Pilot searches. Up to 5 target fragment ions per peptide were automatically selected, and the peak groups were scored following the criteria described in [32]. The peak group confidence threshold was determined based on an FDR analysis using the target-decoy approach, and a 1% extraction FDR threshold was used for all the analyses. Peptides that met the 1% FDR threshold in at least three replicates of a given experimental group were retained, and the peak areas of the target fragment ions of those peptides were extracted across the experiments using an extracted-ion chromatogram window of 4 min. Protein levels were estimated by summing all the transitions from all the peptides for a given protein [33] and normalized to the total intensity at the protein level. Statistical tests were performed in RStudio using the nonparametric Kruskal-Wallis test for multiple hypothesis testing, followed by the post hoc Dunn’s test and adjusting the P values for multiple testing by controlling the FDR using the Benjamini-Hochberg adjustment; proteins were considered altered when an alteration in abundance of at least 20% (fold change of ≤0.83 or ≥1.2) was observed between experimental conditions.

### Bioinformatics analysis

Volcano plots were constructed by plotting fold change values (log2 transformed) on the x-axis and the negative logarithm of the p-value (base 10) on the y-axis using the VolcaNoseR tool (https://goedhart.shinyapps.io/VolcaNoseR/). Gene ontology (GO) enrichment analysis of host proteins with altered abundance upon infection was performed using ShinyGO 0.77 (http://bioinformatics.sdstate.edu/go77/) based on the Biological Process (BP) GO terms. The DAVID Bioinformatics Resources 6.8 (https://david.ncifcrf.gov/home.jsp) was also used to identify proteins according to their GO term [34, 35].

### Western blotting

After thawing, the same amount of protein for each sample was resolved by SDS-PAGE using 12.5% polyacrylamide gels in a Bio-Rad Mini-Protean tetra cell and transferred to a polyvinylidene difluoride membrane at 100 V during 100 min at 4°C. The membranes were blocked for 60 min with 2% BSA in Tris-buffered saline (TBS) containing 0.1% Tween 20 and then incubated at 4°C overnight with primary antibodies. The following primary antibodies were used accordingly: anti-RIG-I (D-12) antibody (sc-376845; Santa Cruz Biotechnology) (1:100), and anti-ISG15 antibody (F-9) (sc-166755; Santa Cruz Biotechnology) (1:100). After several washes with TBS-T (TBS containing 0.1% Tween 20), the membranes were incubated at room temperature with anti-mouse IgG (whole molecule)–peroxidase produced in rabbit (A9044; Sigma) (1:10,000). The membranes were washed again in TBS-T and visualized using NZY supreme ECL HRP substrate (NZYTech) on a VWR Imager. Membranes were also stained using SERVA purple (SERVA electrophoresis; Enzo) to determine the total protein load in each lane according to the manufacturer’s guidelines.

## RESULTS

### Spotted Fever Group *Rickettsia* species display distinct growth dynamics in endothelial cells

We have reported that SFG *Rickettsia* species with different degrees of pathogenicity to humans display completely distinct intracellular fates within the THP-1 macrophages [36, 37]. Specifically, pathogenic *Rickettsia* species subverted macrophage-mediated killing mechanisms, whereas the non-pathogenic *R. montanensis* was rapidly destroyed [36, 37]. In a recent report, Fitzsimmons et al. have also demonstrated distinct survival rates between the non-pathogenic *R. montanensis*, the attenuated *R. rickettsii* strain Iowa, and the highly virulent *R. rickettsii* strain Sheila Smith in primary human endothelial cells [38]. To further explore the intracellular fates between rickettsial species in different cell lines, we infected the hTERT-immortalized human umbilical vein endothelial cell line (HUVEC/TERT2) with different SFG *Rickettsia* species to assess the growth dynamics by immunofluorescence microscopy. We have observed that different SFG *Rickettsia* species display distinct growth dynamics in HUVEC/TERT2 **(Figure 1)**. Of the species herein tested, *R. africae*, the causative agent of African tick-bite fever, presented the fastest growth rates, followed by *R. parkeri*, the causative agent of *R. parkeri* rickettsiosis. On the other hand, *R. massiliae* presented delayed growth dynamics, and no growth was observed for the non-pathogenic *Rickettsia montanensis.* These findings further demonstrate that different SFG *Rickettsia* species interact differently with endothelial cells, likely contributing to explaining the complexity in the spectrum of pathogenesis observed for different *Rickettsia* species.

**Figure 1.**
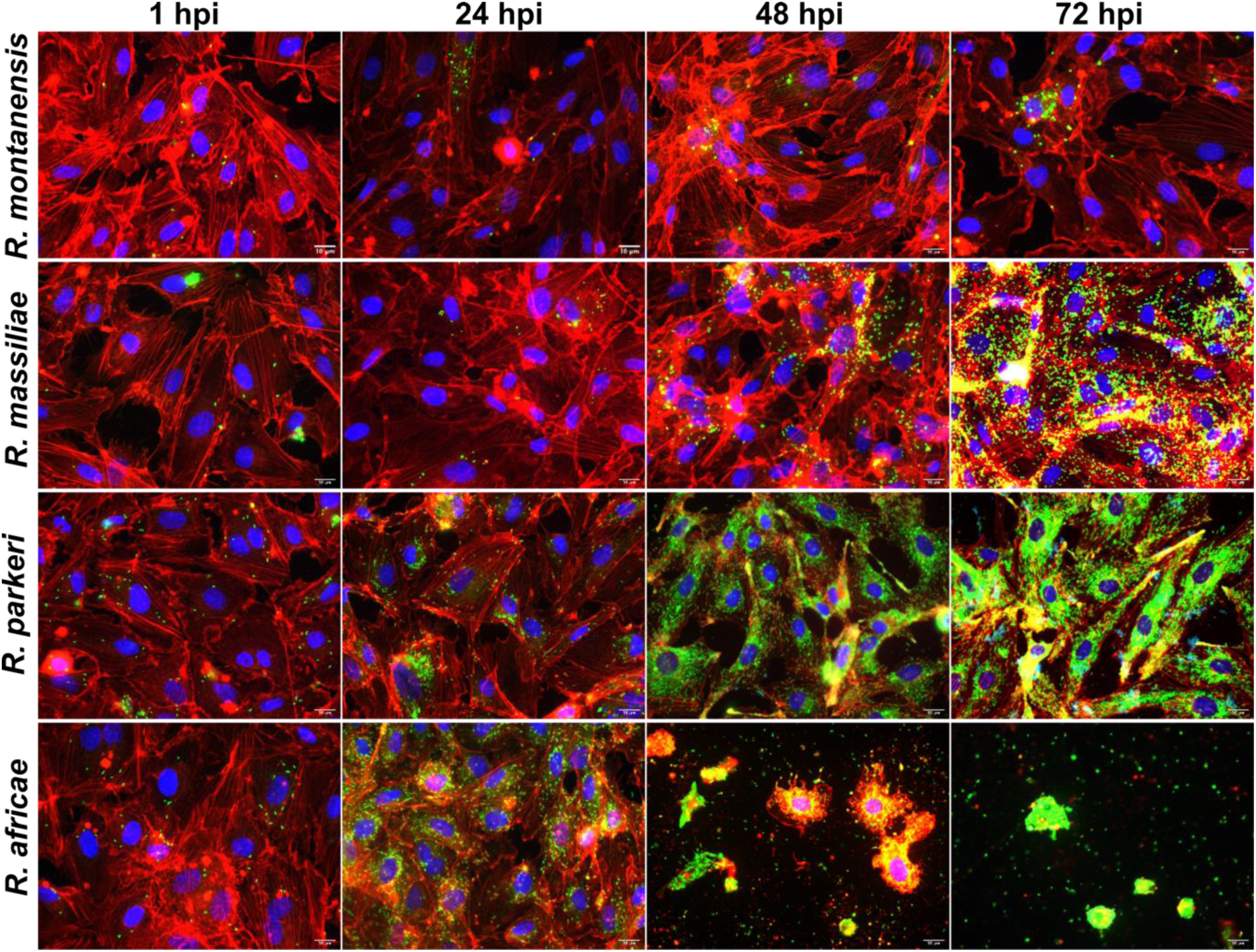
Spotted Fever Group *Rickettsia* species display distinct growth dynamics in endothelial cells. Immunofluorescence microscopy of HUVEC/TERT2 cells infected with *R. montanensis*, *R. massiliae*, *R. parkeri*, and *R. africae* (MOI of 10) at 1 h, 24 h, 48h, and 72 h postinfection. Cells were stained with DAPI (blue) to stain host nuclei, phalloidin (red) to stain actin, and rabbit anti-*Rickettsia* polyclonal antibody NIH/RML I7198 followed by Alexa Fluor 488 (green) to stain *Rickettsia*. Scale bar = 10 μm.

### High-throughput proteomics analysis of endothelial cells infected with different SFG *Rickettsia* species

We have employed a high-throughput proteomics approach to gain further insights into the host signaling pathways and molecular mechanisms elicited by endothelial cells upon infection with SFG *Rickettsia* species. The label-free quantitative proteomics approach (SWATH/DIA-MS/MS) has already been successfully used to profile the dynamics of host-rickettsiae interactions [37, 39]. Hence, total protein extracts from uninfected and *Rickettsia*-infected cells were prepared at 24 hours post-infection (hpi) in a total of 4 biological replicates per experimental condition. The relative protein quantification was performed using liquid chromatography (LC)-SWATH-MS analysis, where a comprehensive library of 2,381 confidently identified proteins was created, and a total of 1,951 proteins were confidently quantified in all samples. Proteins were considered altered when an alteration of at least 20% in abundance (fold change of ≤0.83 or ≥1.2) was observed between uninfected and infected conditions [37, 39]. Using these criteria, significant changes in the content of host proteins were observed upon infection with rickettsial species. Specifically, we found alterations in the content of 536 host proteins (208 enriched and 328 reduced) in *R. africae*-, 351 (219 enriched and 132 reduced) in *R. parkeri*-, 541 (153 enriched and 298 reduced) in *R. massiliae*-, and 182 (83 enriched and 99 reduced) in *R. montanensis*-infected cells **(Figure 2 A-D)**.

**Figure 2.**
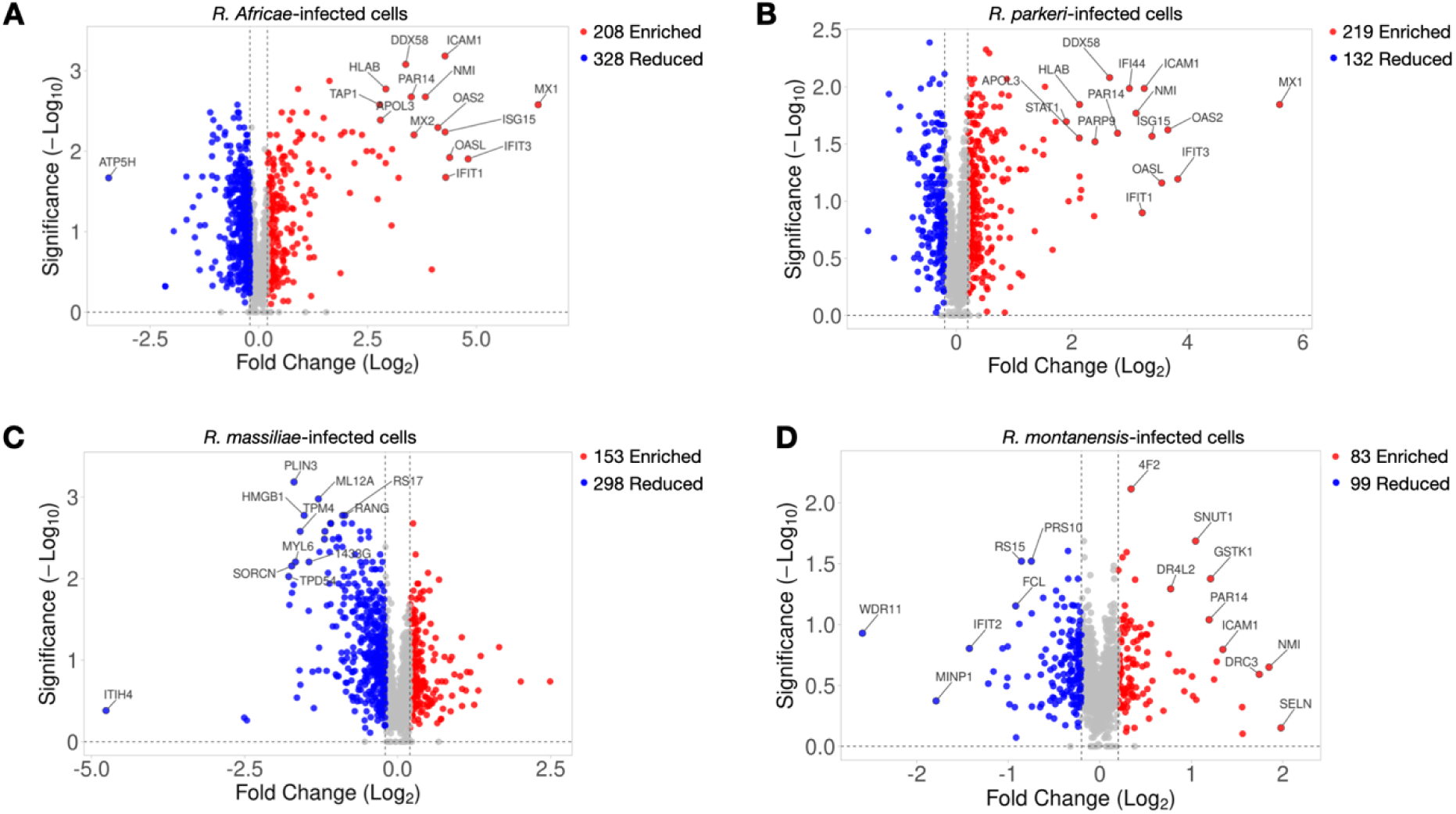
High-throughput proteomics analysis of HUVEC/TERT2 infected with different SFG *Rickettsia* species. Volcano plot representation of changes in protein abundance of HUVEC/TERT2 cells infected with *R. africae* (A), *R. parkeri* (B), *R. massiliae* (C), and *R. montanensis* (D) at 24 hours post-infection, MOI of 10. The 1951 host proteins confidently quantified in all 4 experimental conditions were plotted and considered altered when changes in at least 20% in abundance (fold change of ≤0.83 or ≥1.2) were observed between uninfected and infected conditions. Proteins considered to decrease, not change, or increase their abundance upon infection are represented in blue, gray, and red, respectively. See also Table S1 in the Supplementary Files.

### Gene Ontology analysis of global proteomic alterations in *Rickettsia*-infected endothelial cells

To provide insights into the biological processes modulated by these rickettsial species during infection, we analyzed the host proteins with altered abundance in each infection condition using the ShinyGO enrichment tool [40]. Among the proteins with increased abundance upon infection, we observed an enrichment in several proteins categorized as “viral process” (GO:0016032) in all infection conditions and in “innate immune response” (GO:0045087), “type I interferon signaling pathway” (GO:0060337), and “response to type I interferon” (GO:0034340) in *R. africae*-, R. parkeri-, and *R. massiliae*-infected cells **(Figure 3, left)**. On the other hand, the GO enrichment analysis of proteins with decreased abundance upon infection revealed an enrichment in several proteins categorized as “intracellular transport” (GO:1902582), “RNA fragment catabolic process” (GO:0000292), and “viral processing” (GO:0022415) in *R. africae*-, *R. parkeri*-, and *R. massiliae*-infected cells **(Figure 3, right).** In *R. montanensis*-infected cells, among the proteins with decreased abundance upon infection, we found an enrichment in proteins categorized as “vesicle targeting” (GO:0006903) and “establishment of organelle localization” (GO:0051656).

**Figure 3.**
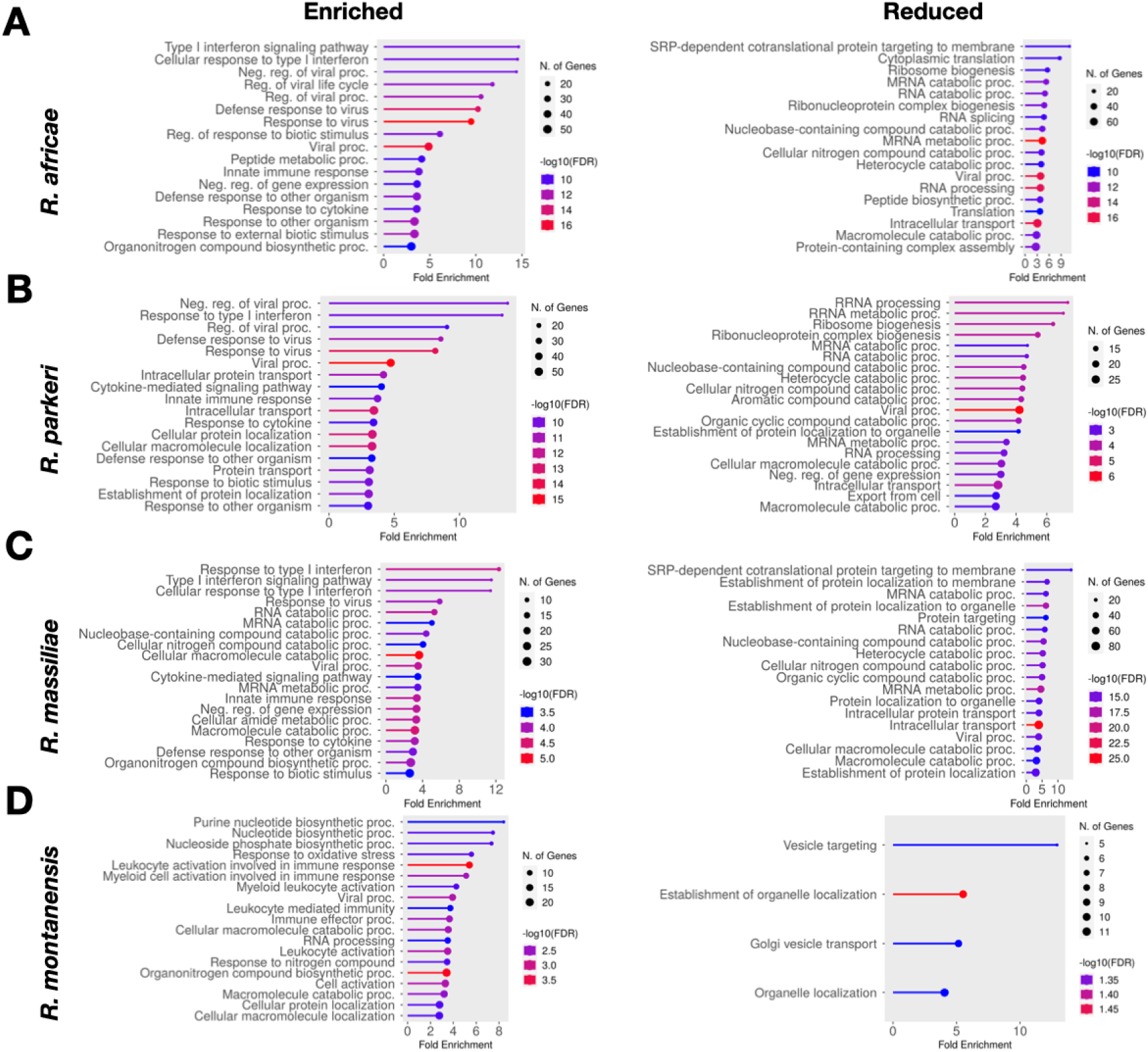
Gene Ontology analysis of global proteomic alterations in *Rickettsia*-infected endothelial cells. GO enrichment analysis of host proteins with altered abundance (enriched – left and reduced – right) upon infection was performed using ShinyGO 0.77 (http://bioinformatics.sdstate.edu/go77/) based on the Biological Process (BP) GO terms. Analysis of the host proteins with altered abundance upon infection with *R. africae* (A), *R. parkeri* (B), *R. massiliae* (C), and *R. montanensis* (D). The dot size represents the number of host proteins with altered abundance for the specific infection condition of each GO term. The dot color indicates the false discovery rate (FDR) as p-values, and the X-axis denotes the enrichment factor of altered proteins.

### Type I interferon response in endothelial cells infected with different SFG *Rickettsia* species

Endothelial cells are key players in the innate immune response to pathogenic insults and are among the first line of defense against rickettsial infections [8, 10, 13, 21–23]. To gain further insights into the innate immune response elicited by endothelial cells upon infection with different *Rickettsia* species, we compared alterations in the abundance of individual proteins categorized as GO:0045087-“innate immune response” in *Rickettsia*-infected cells **(Figure 4A)**. Of note, we found an increased abundance of several proteins involved in innate immunity, including the antiviral innate immune response receptor RIG-I (DDX58; O95786), the protein mono-ADP-ribosyltransferase PARP9 (PARP9; Q8IXQ6), and the protein mono-ADP-ribosyltransferase PARP14 (PAR14; Q460N5), among others. This increase was more pronounced in *R. africae*- and *R. parkeri*-infected cells compared to *R. massiliae*- and *R. montanensis*-infected cells, correlating well with the number of bacteria present within these cells at this time post-infection. Among these proteins, we have also found several that were categorized as GO:00600337 – “Type I Interferon” **(Figure 4B)**, including several interferon-induced proteins such as interferon-induced protein with tetratricopeptide repeats 1 (IFIT1; P09914), 2 (IFIT2; P09913), 3 (IFIT3; O14879), interferon-induced GTP-binding protein Mx1 (MX1; P20591) and 2 (MX2; P20592), and the interferon-induced 35 kDa protein (IFI35; P80217). An increased abundance of several 2’-5’-oligoadenylate synthase proteins, such as the 2’-5’-oligoadenylate synthase-like protein (OASL; Q15646), the 2’-5’-oligoadenylate synthase 2 (OAS2; P29728), and the 2’-5’-oligoadenylate synthase 3 (OAS3; Q9Y6K5) was also observed in *R. africae*- and *R. parkeri*-infected cells at this time point. To further validate these results, we performed a WB analysis of uninfected and *Rickettsia*-infected cells, at 24 hpi, for the DDX58/RIG-I and the ISG15 proteins **(Figure 4C, D)**. These results confirmed an increase in DDX58/RIG-I and ISG-15 abundance in *R. africae*- and *R. parkeri*-infected cells, more pronounced in the former.

**Figure 4.**
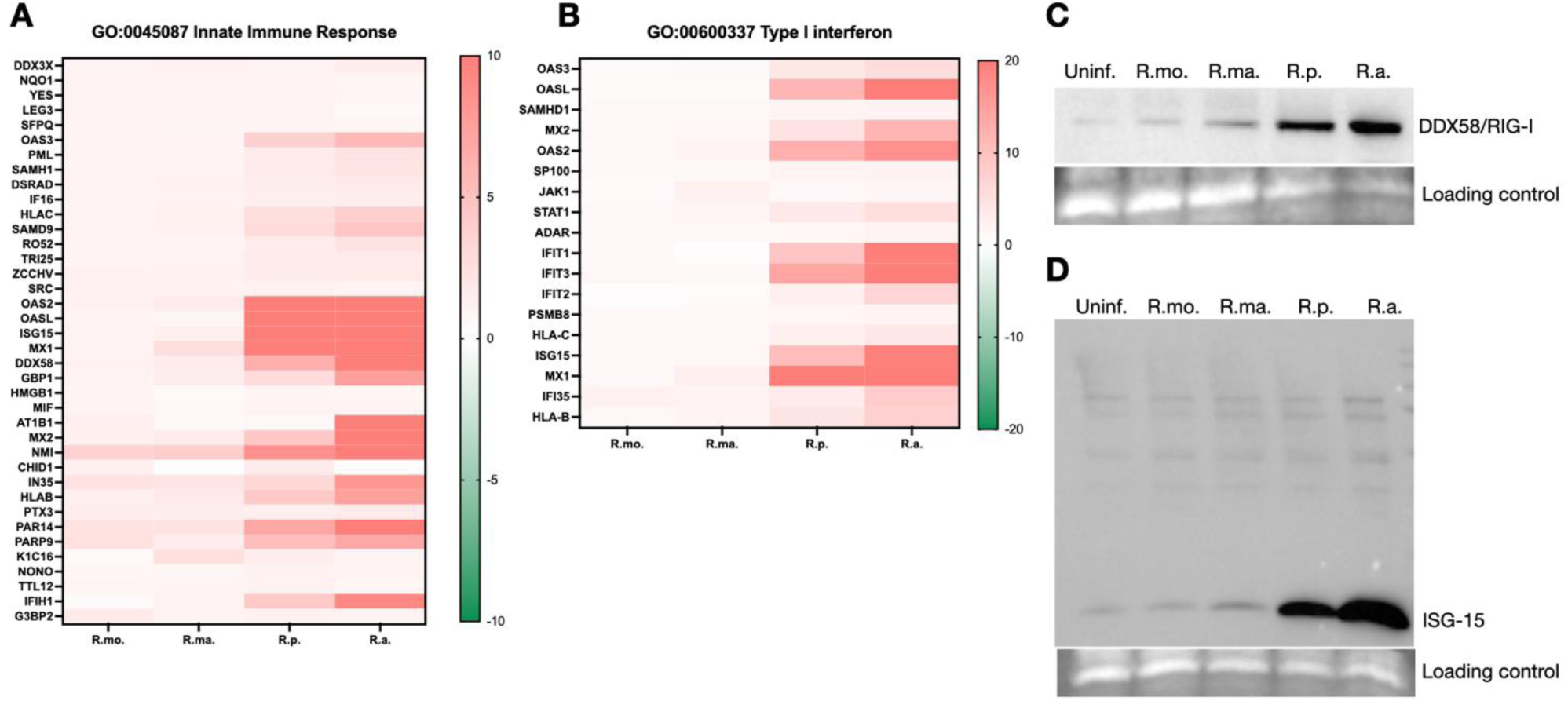
Type I interferon response in endothelial cells infected with different SFG *Rickettsia* species. (A-B) Heatmap representation of host proteins categorized as Innate Immune Response - GO:0045087 (A) and Type I Interferon – GO:00600337 (B) in endothelial cells infected with different SFG *Rickettsia* species. The heatmaps compare alterations between infected and uninfected conditions (*R. montanensis*-infected vs. uninfected – R.mo.; *R. massiliae*-infected vs. uninfected – R.ma.; *R. parkeri*-infected vs. uninfected – R.p.; *R. africae*-infected vs. uninfected – R.a.). The green (decreased) and red (increased) color scales indicate alterations in protein abundance (log_2_ ratio). (C-D) Western blot analysis of total protein extracts (15 µg) from uninfected THP-1 macrophages (uninf.) and THP-1 macrophages infected with *R. montanensis* (R.mo.), *R. parkeri* (R.p.), *R. africae* (R.a.), and *R. massiliae* (R.ma.). Protein samples were probed for retinoic acid-inducible gene I (RIG-1 [accession no. O95786]), and ubiquitin-like protein ISG15 (accession no. P05161), and immunoblot analysis with SERVA purple was used as the protein loading control. See also Table S2 in the Supplementary Files.

### *Rickettsia africae* induces cell type-specific and common host responses during infection

We have previously profiled the alterations in the host proteome of THP-1 macrophages upon infection with different SFG *Rickettsia* species [37]. Since 1,321 host proteins were commonly quantified in our previous and current datasets, we searched for host responses common to both *Rickettsia*-infected endothelial cells and macrophages, as well as those specific to each cell type. Using *R. africae* as our model organism, we built a correlation plot of the 1,321 host proteins and their respective log2 fold change values observed at 24 hpi in both cell types **(Figure 5A)**. Of note, we found 88 host proteins that commonly changed their abundance upon infection with *R. africae*. Of these 88 proteins, 48 showed increased abundance, whereas 40 proteins were found in lower abundance in both cell types **(Figure 5B, C)**. Gene ontology analysis of the 48 proteins that showed increased abundance during *R. africae* infection in both cell types demonstrated an enrichment in proteins categorized as “type I interferon signaling pathway” (GO:0060337), suggesting that both cell types contribute to type I interferon responses during rickettsial infections **(Figure 5D, and Table S3)**. On the other hand, gene ontology analysis of the 40 proteins found in lower abundance in both cell types showed an enrichment in proteins categorized as “positive regulation of pinocytosis” (GO:0048549), “nuclear division” (GO:0000280), “neutrophil mediated immunity” (GO:0002446), and “granulocyte activation” (GO:0036230) **(Figure 5E, and Table S3)**. We have also searched for proteins that altered abundance in a cell-type-specific manner and found that 11 host proteins increase abundance in HUVEC/TERT2-infected cells but decrease in THP-1-infected cells, and 20 proteins decrease abundance in HUVEC/TERT2-infected cells but increase in THP-1-infected cells **(Figure 5F)**. Among these proteins is the sodium/potassium-transporting ATPase subunit beta-1 (ATP1B1; P05026), which has already been described to play a role in innate immunity [41], and the myristoylated alanine-rich C-kinase substrate (MARCKS; P29966) that promotes the migration and adhesion on inflammatory cells and secretion of cytokines such as tumor necrosis factor (TNF) during inflammation [42].

**Figure 5.**
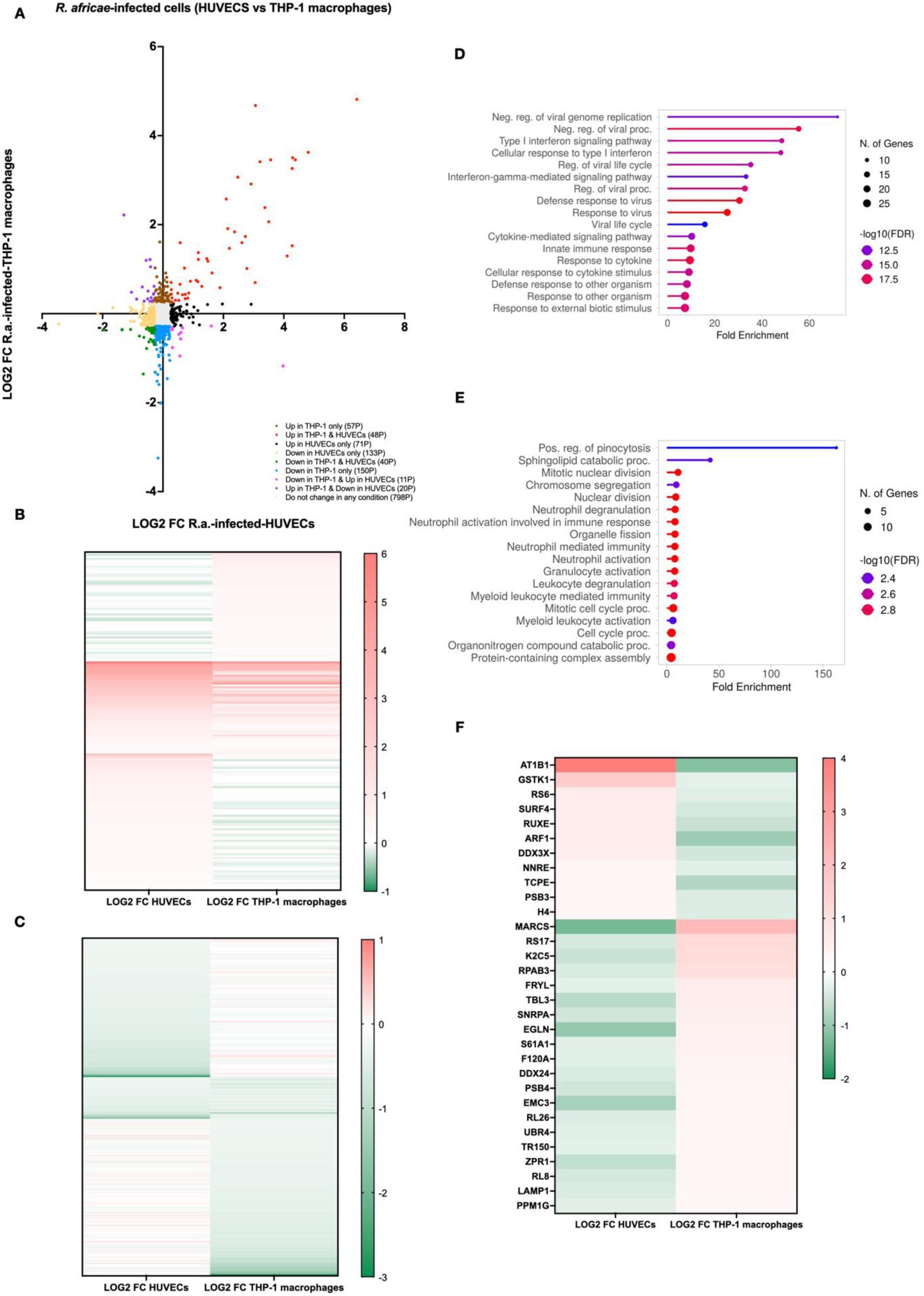
Cell-type-specific and common host responses to infection with *R. africae*. (A) Correlation plot of the 1321 host proteins confidently quantified in *Rickettsia*-infected THP-1 macrophages and HUVEC/TERT2 cells. Individual proteins were plotted according to their log2 fold change values in *R. africae*-infected HUVEC/TERT2 cells (x-axis) and the corresponding log2 fold change values in *R. africae*-infected THP-1 macrophages (y-axis) at 24hpi. Color code depicts proteins that do not change abundance in any infection condition (gray; 798 proteins), increase abundance only in THP-1-infected cells (brown; 57 proteins), increase abundance in both THP-1- and HUVEC/TERT2-infected cells (red; 48 proteins), increase abundance in HUVEC/TERT2-infected cells only (black; 71 proteins), decrease abundance in HUVEC/TERT2-infected cells only (yellow; 133 proteins), decrease abundance in both THP-1- and HUVEC/TERT2-infected cells (green; 40 proteins), decrease abundance in THP-1-infected cells only (blue; 150 proteins), increase abundance in HUVEC/TERT2-infected cells and decrease in THP-1-infected cells (pink; 11 proteins), decrease abundance in HUVEC/TERT2-infected cells and increase in THP-1-infected cells (purple; 20 proteins). (B) Heatmaps depicting the log2 fold change values for the 57 proteins that increase abundance only in THP-1-infected cells, the 48 proteins that increase abundance in both THP-1- and HUVEC/TERT2-infected cells, and the 71 proteins that increase abundance in HUVEC/TERT2-infected cells only. (C) Heatmaps depicting the log2 fold change values for the 133 proteins that decrease abundance in HUVEC/TERT2-infected cells only, the 40 proteins that decrease abundance in both THP-1- and HUVEC/TERT2-infected cells, and the 150 proteins that decrease abundance in THP-1-infected cells only. (D-E) GO enrichment analysis of host proteins that commonly changed abundance, enriched – (D) and reduced – (E), upon infection in macrophages and endothelial cells with *R. africae*. GO analysis was performed using ShinyGO 0.77 (http://bioinformatics.sdstate.edu/go77/) based on the Biological Process (BP) GO terms. The dot color indicates the false discovery rate (FDR) as p-values, and the X-axis denotes the enrichment factor of altered proteins. (F) Heatmaps depicting the log2 fold change values for the 11 proteins that increase abundance in HUVEC/TERT2-infected cells and decrease in THP-1-infected cells and the 20 proteins that decrease abundance in HUVEC/TERT2-infected cells and increase in THP-1-infected cells. See also Table S3 in the Supplementary Files.

## DISCUSSION

Endothelial infection is a hallmark of SFG *Rickettsia* pathogenesis, yet the dynamics of host-pathogen interactions across species with differing virulence profiles have remained poorly defined. In this study, we leveraged high-throughput quantitative proteomics using HUVEC/TERT2 cells to dissect the host cellular response to infection with four SFG *Rickettsia* species, providing a comparative analysis of their distinct growth dynamics and the associated cellular pathways impacted within endothelial cells.

Consistent with their virulence profiles, *R. africae*, followed by *R. parkeri*, demonstrated higher replication rates within HUVEC/TERT2 cells, whereas *R. massiliae* exhibited delayed growth and *R. montanensis* failed to replicate effectively in these cells. These differences in growth dynamics between pathogenic and non-pathogenic species mirror our previous observations in THP-1 macrophages [37] and support the paradigm that more pathogenic species are skilled at subverting host cellular defenses, allowing robust intracellular replication. By extending this observation to endothelial cells, our results emphasize the central role of cellular context in determining the success and nature of the infection. These conclusions are also supported by the work of Fitzsimmons et al., who demonstrated that only the highly virulent *R. rickettsii* Sheila Smith strain efficiently replicated in primary human dermal microvascular endothelial cells (HDMECs), while the attenuated *R. rickettsii* Iowa strain showed minimal replication, and *R. montanensis* failed to replicate [38].

Our proteomic analyses revealed both common and distinct host responses elicited by the four species. Gene Ontology analyses uncovered distinct functional enrichments associated with each pathogen. However, all pathogenic species triggered enrichment of host proteins linked to “viral process”, “type I interferon signaling,” and “response to type I interferon”, highlighting the central role of these pathways in the host response. Conversely, downregulated protein sets in pathogen-infected cells were enriched in categories related to mRNA/RNA catabolic/metabolic processing (interestingly also with increased accumulation in *R. massiliae*-infected cells) or protein translation/targeting (co-translational targeting to membrane). These findings anticipate a perturbation of host transcription and translation machineries during endothelial infection with mildly pathogenic rickettsiae and call for further investigation into how such modulation may influence bacterial replication and contribute to rickettsial pathogenesis. *R. montanensis*-infected endothelial cells mostly exhibited decreased abundance of “vesicle targeting” and “organelle localization” proteins, suggesting a distinct, potentially abortive infection process.

Notably, an increased abundance of host proteins associated with antiviral immunity, including RIG-I (DDX58), ISG15, IFITs, MX1, MX2, IFI35, and OAS family members, was observed across species, with the most robust induction in *R. africae*- and *R. parkeri*-infected endothelial cells. These results highlight the pivotal role of the type I interferon pathway in endothelial defense against SFG *Rickettsia*, aligning with prior observations in THP-1 macrophages for the same pathogenic species [37], and underscoring the role of this pathway as a core component of host defense. The increased accumulation of the 2′-5′-oligoadenylate synthetase (OAS) family of innate immune sensors (OASL, OAS2, and OAS3) - which detect cytosolic double-stranded RNA (dsRNA) and activate the latent RNase L enzyme, leading to the degradation of both pathogen-derived and host RNAs and thereby inhibiting pathogen replication [43] - is consistent with our observations in THP-1 macrophages [37]. Intriguingly, although critical in antiviral immunity, there are only a few reports of their involvement in bacterial infections, and mechanistic details remain largely unknown [44–46]. This raises important questions about the functional consequences of their increased accumulation and the nature of the rickettsial/host ligands being recognized. Similarly, the accumulation of RIG-I suggests potential cytosolic detection of RNA, again prompting important questions about the mechanisms underlying the interaction between *Rickettsia* and these critical immune sensors. As with OASL proteins, the interferon-stimulated GTPases MX1 and MX2 are primarily known for their antiviral activity, which evolved to target different stages of viral replication in different types of viruses [47, 48]. To our knowledge, their role in bacterial infections has been suggested to be rather limited and mostly indirect, likely reflecting bystander IFN signaling [49]. Therefore, the increased accumulation observed upon *R. africae* and *R. parkeri* infection urges further research to fully explore potential, yet undiscovered, roles of MX1 and MX2 in antirickettsial immunity. Indeed, several non-antiviral activities have been reported for MX2, such as the regulation of mitochondrial metabolism or inhibition of nuclear import of non-viral cargo [50]. While the full functional consequences of these non-antiviral activities remain unknown [50], we can hypothesize that MX proteins may contribute to host defense against intracellular bacteria like *Rickettsia* through novel, cell-type-specific mechanisms that extend beyond classical antiviral responses. Likewise, IFIT1-3 are mostly known for their antiviral activity through binding to non-self RNAs, inhibiting their translation or replication, as well as through interactions with cellular components, including modulation of the MAVS signaling axis and apoptosis [51, 52]. While their functions in viral infections are well established, evidence for direct roles in bacterial infections remains limited [53–56]. However, recent findings in *Mycobacterium tuberculosis* suggest that IFITs may also contribute to antibacterial immunity, with elevated IFIT expression directly reducing bacterial burden [57]. Given their robust induction during rickettsial infection, a deeper understanding of IFIT functions in this context is warranted to uncover potential, yet unrecognized, roles in host defense against obligate intracellular bacteria.

Interestingly, the infection-induced accumulation of ISG15 followed a gradient - *R. africae* > *R. parkeri* > *R. massiliae* > *R. montanensis* - revealing a distinct pattern from that previously observed in THP-1 macrophages [37], where comparable levels of free ISG15 were induced upon infection with the three mildly pathogenic species. Whether this discrepancy reflects differences in cell-type-specific interferon responses or a greater tropism and replication capacity of *R. massiliae* in macrophages remains to be determined. While ISG15 expression was clearly induced, global ISGylation of host proteins remained low under all endothelial infection conditions. This contrasts with previous findings in THP-1 macrophages, where ISGylation was detectable upon infection with the same pathogenic species [37]. These results suggest that rickettsial species may differentially regulate post-translational modifications such as ISGylation depending on the host cell type. Remarkably, ISG15 and ISGylation have been reported to increase in human microvascular endothelial cells infected with the highly virulent *R. conorii* [58]. The low ISGylation levels observed in our study may reflect both pathogen- and host cell-specific regulatory mechanisms. Further studies are, therefore, required to identify the host and/or bacterial proteins that undergo ISGylation (or deISGylation) during rickettsial infection across different cell types and to elucidate how the regulation of this post-translational modification influences the replication and pathogenic potential of distinct *Rickettsia* species.

In parallel, direct comparison between endothelial and THP-1 macrophage responses revealed a core signature of 88 host proteins with concordant abundance changes upon *R. africae* infection, many of which are components of the type I interferon pathway. These commonalities point to fundamental host-pathogen interactions that may define the early cellular response to rickettsial invasion, regardless of cellular context. Yet, the identification of cell type-specific responses, such as the opposing regulation of ATP1B1 and MARCKS between HUVEC/TERT2 and THP-1, highlights the complex interplay between pathogen and host, suggesting that the cellular environment may significantly shape the trajectory and outcome of infection.

Taken together, this study provides a comprehensive view of the proteomic landscape of endothelial infection by SFG *Rickettsia* species, revealing both shared and species-specific host responses that align with their distinct pathogenic profiles. The strong induction of antiviral ISGs - typically associated with viral infections - suggests that the host perceives *Rickettsia* through pathways normally reserved for antiviral defense, potentially via cytosolic sensing of rickettsial or host-derived nucleic acids. This highlights the need to uncover how these sensors influence bacterial clearance, immune modulation, and pathogenesis. These findings deepen our understanding of the molecular mechanisms underlying SFG *Rickettsia* host specificity and virulence and identify immune signatures that may be used to predict the pathogenic potential of emerging or poorly characterized *Rickettsia* species.

## CONTRIBUTIONS

P.C., I.S. conceptualized experiments. P.C., C.S., conducted the investigation and analyzed the data. I.S., B.M., provided resources. P.C. acquired the funding. P.C., I.S., contributed to the original draft of the manuscript. All authors discussed the results, reviewed, and edited the manuscript.

## ACKNOWLEDGMENTS

This work was funded through the COMPETE 2020 - Operational Programme for Competitiveness and Internationalisation and Portuguese national funds via FCT – Fundação para a Ciência e a Tecnologia, under project 2022.02702.PTDC (TalkRick) (https://doi.org/10.54499/2022.02702.PTDC) and UIDB/04539/2020, UIDP/04539/2020 and LA/P/0058/2020, and the National Mass Spectrometry Network (POCI-01–0145-FEDER-402–022125 Ref. ROTEIRO/0028/2013). IS thanks start-up funding from Old Dominion University. PC thanks funding from the European Union’s Horizon 2020 research and innovation programme under grant agreement No. 951921.

